## Supplemental Tables for "A Conserved Switch Controls Virulence, Sporulation, and Motility in *C. difficile*"

**Table S1.** Bacterial Strains and plasmids

| Plasmid or Strain | | Relevant genotype or features | Source, construction or reference |
|---|---|---|---|
| **Strains** | | | |
| ***E. coli*** | | | |
| | DH5α | | |
| | HB101 | F$^-$ *mcrB mrr hsdS20*($r_B^-$ $m_B^-$) *recA13 leuB6 ara-14 proA2 lacY1 galK2 xyl-5 mtl-1 rpsL20* (conjugation) | B. Dupuy |
| ***C. difficile*** | | | |
| | 630Δ*erm* | Erm$^S$ derivative of strain 630 | (*53*) |
| | RT1075 | 630Δ*erm sigD*::*erm* | (*54*) |
| | MC324 | 630Δ*erm* pMC123 | (*55*) |
| | MC855 | 630Δ*erm spo0A*::*erm* pMC123 | (*22*) |
| | MC1003 | 630Δ*erm spo0A*::*erm* pMC674 | This study |
| | MC1615 | 630Δ*erm spo0E*::*erm* | This study |
| | MC1698 | 630Δ*erm spo0E*::*erm* pMC980 | This study |
| | MC1699 | 630Δ*erm spo0E*::*erm* pMC123 | This study |
| | MC1968 | 630Δ*erm spo0E*::*erm* pMC1093 | This study |
| *B. subtilis* | | | |
| | 1A1 | strain 168 lineage | (*56*) |
| | 1S143 | MF2339; PY79 *spo0A::kan* | M. Fujita/BGSC |
| | BKE13640 | Δ*spo0E::erm* | (*56*) |
| | MC2261 | 1S143→1A1; *spo0A::kan* | This study |
| | MC2400 | BKE13640→1A1; Δ*spo0E::erm* | This study |
| **Plasmids** | | | |
| | pBB1364 | *kan* | (*57*) |
| | pCE240 | *C. difficile* TargeTron® construct based on pJIR750ai (group II intron, *ermB::*RAM, *ltrA*); *catP* | C. Ellermeier |
| | pMC123 | *E. coli-C. difficile* shuttle vector; *bla, catP* | (*58*) |
| | pMC228 | pMC123 with *spo0E*-targeted intron | This study |
| | pMC674 | pMC123 s*po0A::*3xFLAG | (*22*) |
| | pMC980 | pMC123 *CD3272-spo0E* | This study |
| | pMC1093 | pMC123 *spo0E::*3xFLAG | This study |

**Table S2.** Filtered proteins identified in Spo0E-FLAG co-immunoprecipitation

| Gene Locus | -Log P value[a] | $Log_2$ Intensity/control[b] |
|---|---|---|
| CD630_12140 (Spo0A) | 8.0 | 1.4 |
| CD630_24670 (LepA) | 7.0 | 1.3 |
| CD630_32710 (Spo0E) | 6.7 | 1.7 |
| CD630_36680 (RstA) | 5.5 | 1.3 |
| CD630_21730 | 5.4 | 1.4 |
| CD630_20070 | 5.3 | 1.4 |
| CD630_29560 | 5.2 | 1.3 |
| CD630_35120 | 5.0 | 1.4 |
| CD630_21230 | 4.5 | 1.3 |
| CD630_03400 | 4.4 | 1.2 |
| CD630_29800 | 4.2 | 1.2 |
| CD630_00511 | 4.2 | 1.9 |
| CD630_08210 | 4.0 | 1.3 |
| CD630_22640 | 3.7 | 1.2 |
| CD630_26460 | 3.2 | 1.3 |
| CD630_34700 | 3.1 | 1.2 |
| CD630_22090 | 2.6 | 1.4 |
| CD630_03410 | 2.6 | 1.2 |
| CD630_05590 | 2.1 | 1.2 |

[a]Negative log of *t*-test between average protein intensities of Spo0E-FLAG and negative control pulldown
[b]Ratio of averaged $log_2$ transformed intensities between Spo0E-FLAG and negative control pulldown

**Table S3.** Filtered proteins identified in Spo0A-FLAG co-immunoprecipitation

| Gene locus | -Log P value[a] | $Log_2$ Intensity/control[b] |
|---|---|---|
| CD630_15790 (PtpC) | 5.5 | 1.5 |
| CD630_12140 (Spo0A) | 5.5 | 1.3 |
| CD630_32710 (Spo0E) | 5.3 | 1.5 |
| CD630_12310 | 5.2 | 1.2 |
| CD630_03410 | 4.9 | 1.2 |
| CD630_35230 | 4.8 | 1.3 |
| CD630_24040 | 4.7 | 1.2 |
| CD630_21230 | 4.6 | 1.3 |
| CD630_P10 | 3.8 | 1.4 |
| CD630_20070 | 3.8 | 1.4 |
| CD630_19320 | 3.8 | 1.2 |
| CD630_13060 | 3.7 | 1.3 |
| CD630_25220 | 3.6 | 1.3 |
| CD630_23980 | 3.6 | 1.3 |
| CD630_05230 | 3.5 | 1.2 |
| CD630_00200 | 3.3 | 1.2 |
| CD630_01500 | 3.2 | 1.2 |
| CD630_35940 | 3.0 | 1.2 |
| CD630_18490 | 2.9 | 1.3 |
| CD630_35460 | 2.9 | 1.2 |
| CD630_19670 | 2.6 | 1.2 |
| CD630_19640 | 2.4 | 1.2 |
| CD630_21800 | 2.4 | 1.2 |
| CD630_12470 | 2.4 | 1.2 |
| CD630_19660 | 1.8 | 1.2 |

[a]Negative log of *t*-test between average protein intensities of Spo0A-FLAG and negative control pulldown
[b]Ratio of averaged $log_2$ transformed intensities between Spo0A-FLAG and negative control pulldown

**Table S4.** Oligonucleotides

| Primer | Sequence (5'→3')[a] | Use/locus tag/reference |
|---|---|---|
| oMC513 | GCGGATCCGACAAAATATAATATTGTTTGATAAAATG | Forward primer for *CD630_32710* |
| oMC514 | GACGGATCCCTGTGGGCTATTTGCTTAGG | Reverse primer for *CD630_32710* |
| oMC515 | AAAAGCTTTTGCAACCCACGTCGATCGTGAA-AACTCTTCTTGA-GTGCGCCCAGATAGGGTG | *CD630_32710* IBS targeting *spo0E* |
| oMC516 | CAGATTGTACAAATGTGGTGATAACAGATAAGTC-TCTTGAAA-TAACTTACCTTTCTTTGT | *CD630_32710* EBS1 targeting *spo0E* |
| oMC517 | CGCAAGTTTCTAATTTCGGTT-GAGTT-TCGATAGAGGAAAGTGTCT | *CD630_32710* EBS2 targeting *spo0E* |
| oMC2589 | AGGGATCCATCACTAAAATTGTAACAAGTATGATAC | Forward primer for *CD630_32720-32710* complement |
| oMC2590 | CAGGAATTCACAGATAATTTACACATCAGAAATAC | Reverse primer for *CD630_32720-32710* complement |
| oMC3546 | GCAGGATCCGAATCCAAGACCGAGGATTACG | Forward primer upstream of *BSU1364* |
| oMC3547 | CAGGAATTCGTTCCTTCATTAGTTCCCTCTTTC | Reverse primer downstream of *BSU1364* |

[a]Restriction sites underlined

**Table S5.** Vector and strain construction

| Vector/Strain | Construction details |
|---|---|
| pMC228 | The group II intron of pCE240 was targeted to *spo0E* at nucleotide 75 by splicing PCR using oMC515, oMC516, oMC517, and EBSu as outlined in the TargeTron user manual (Sigma-Aldrich). The group II *CD1558*-targeted intron was subcloned using the BsrGI and HindIII sites into pCE240. A 5.45 kb SphI/SfoI fragment was then cloned as SphI/SnaBI into pMC123. |
| pMC980 | A 963 bp product containing full-length operon *CD630_32720*-*spo0E* to complement the *spo0E* mutant was generated using primers oMC2589 and oMC2590 and cloned into pMC123 using BamHI and EcoRI sites. |
| pMC1093 | A 683 bp fragment encoding *C. difficile spo0E* with a C-terminal 3xFLAG tag driven by the *spo0E* native promoter was synthesized by Genscript and cloned into pMC123 using BamHI and EcoRI sites. |
| pMC1258 | A 792 bp fragment containing *B. subtilis spo0E (BSU13640)* and upstream region from strain 1A1 was amplified with primers oMC3546 and oMC3547, and cloned as BamHI/EcoRI into pMC123. |
| pMC1259 | The 963 bp fragment containing *C. difficile spo0E* and upstream region from pMC980 was cloned using BamHI and EcoRI sites into pBB1364 for integration at the *B. subtilis sacA* locus. |
