## Supplemental Figures for "A Conserved Switch Controls Virulence, Sporulation, and Motility in *C. difficile*"

### Slide 1
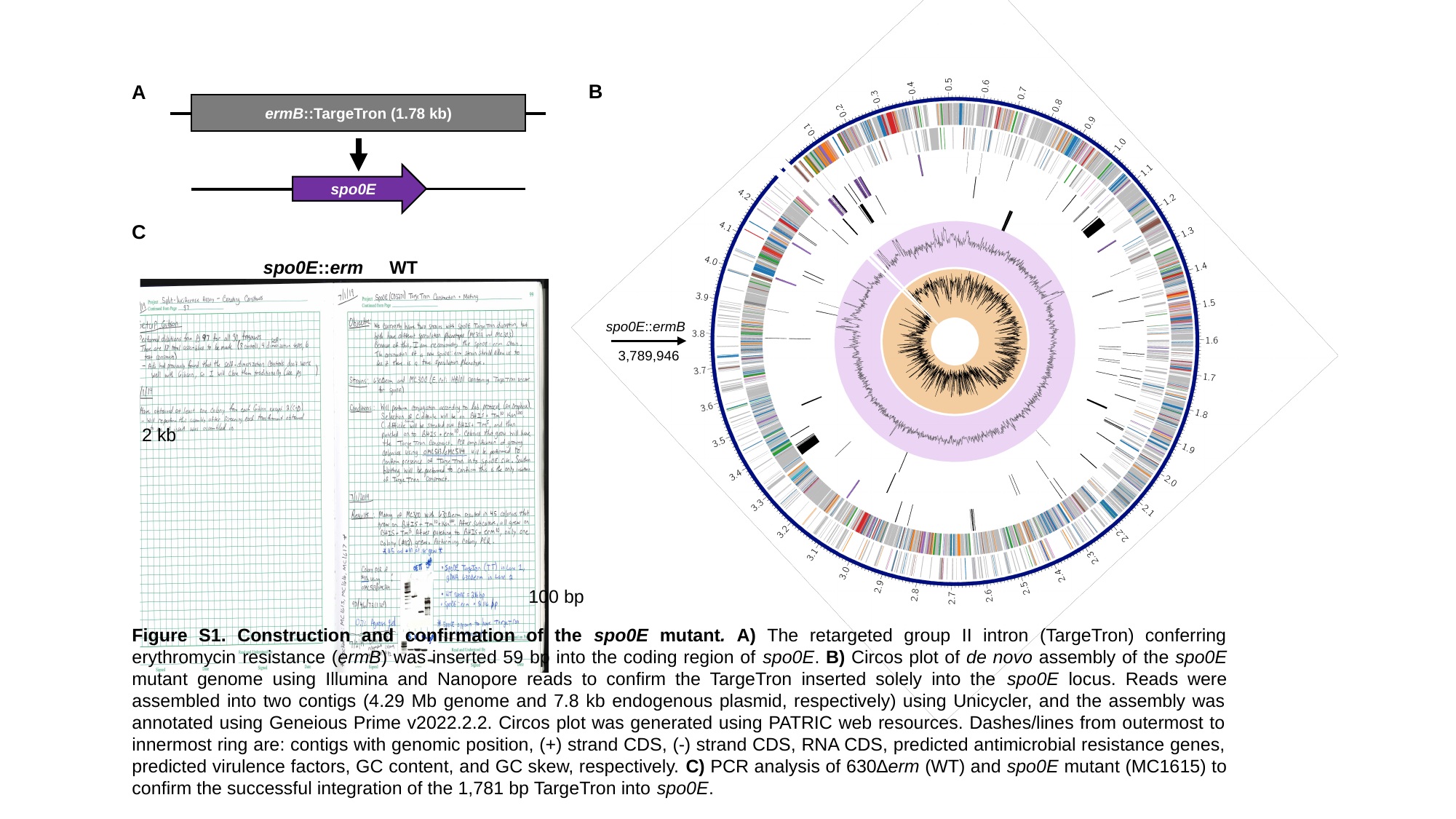

spo0E::ermB
3,789,946
B
A
ermB::TargeTron (1.78 kb)
spo0E
C
spo0E::erm
WT
2 kb
100 bp
Figure S1. Construction and confirmation of the spo0E mutant. A) The retargeted group II intron (TargeTron) conferring erythromycin resistance (ermB) was inserted 59 bp into the coding region of spo0E. B) Circos plot of de novo assembly of the spo0E mutant genome using Illumina and Nanopore reads to confirm the TargeTron inserted solely into the spo0E locus. Reads were assembled into two contigs (4.29 Mb genome and 7.8 kb endogenous plasmid, respectively) using Unicycler, and the assembly was annotated using Geneious Prime v2022.2.2. Circos plot was generated using PATRIC web resources. Dashes/lines from outermost to innermost ring are: contigs with genomic position, (+) strand CDS, (-) strand CDS, RNA CDS, predicted antimicrobial resistance genes, predicted virulence factors, GC content, and GC skew, respectively. C) PCR analysis of 630∆erm (WT) and spo0E mutant (MC1615) to confirm the successful integration of the 1,781 bp TargeTron into spo0E.

### Slide 2
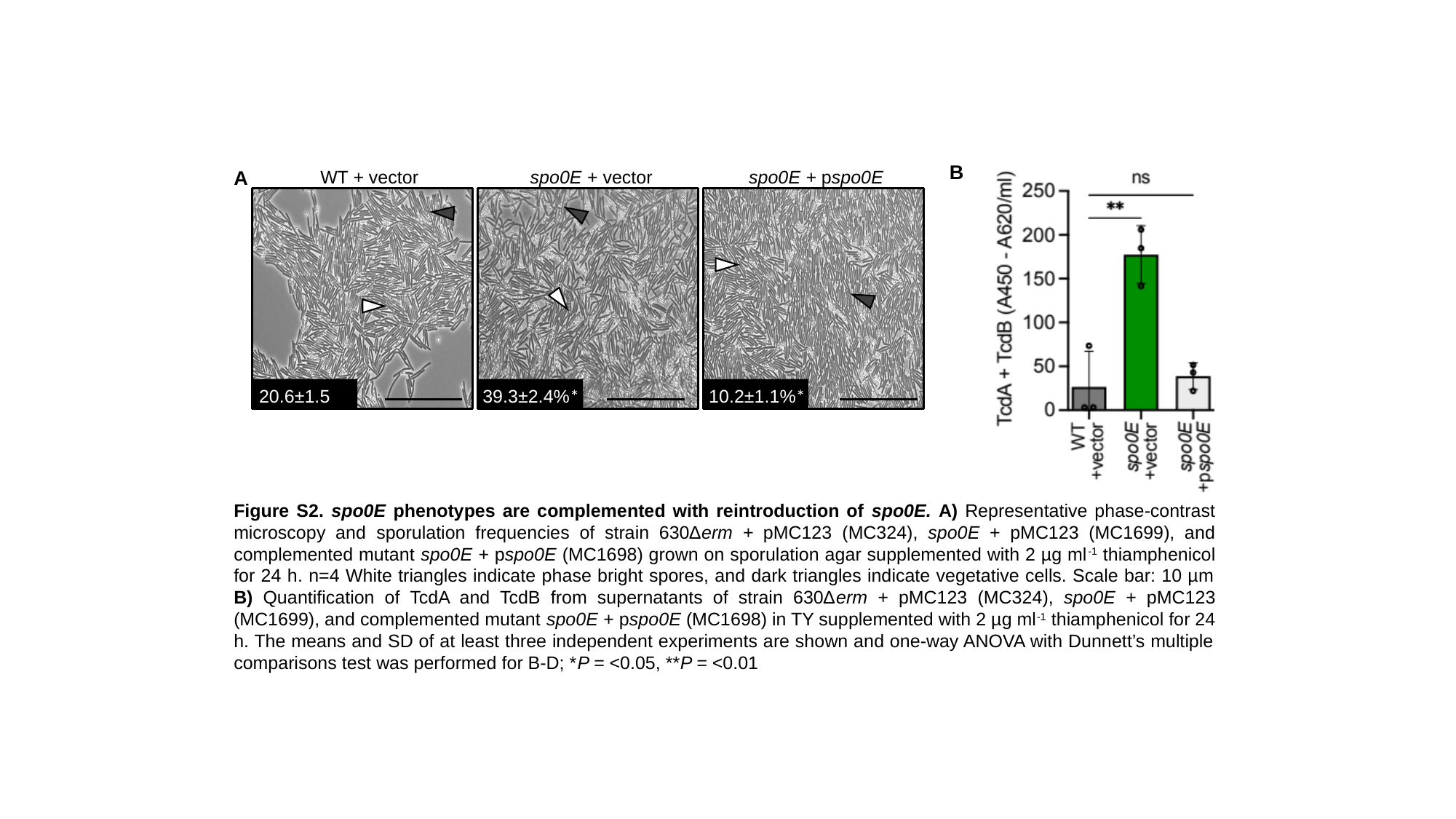

B
A
WT + vector
spo0E + vector
spo0E + pspo0E
20.6±1.5%
39.3±2.4%*
10.2±1.1%*
Figure S2. spo0E phenotypes are complemented with reintroduction of spo0E. A) Representative phase-contrast microscopy and sporulation frequencies of strain 630∆erm + pMC123 (MC324), spo0E + pMC123 (MC1699), and complemented mutant spo0E + pspo0E (MC1698) grown on sporulation agar supplemented with 2 µg ml^-1^ thiamphenicol for 24 h. n=4 White triangles indicate phase bright spores, and dark triangles indicate vegetative cells. Scale bar: 10 µm B) Quantification of TcdA and TcdB from supernatants of strain 630∆erm + pMC123 (MC324), spo0E + pMC123 (MC1699), and complemented mutant spo0E + pspo0E (MC1698) in TY supplemented with 2 µg ml^-1^ thiamphenicol for 24 h. The means and SD of at least three independent experiments are shown and one-way ANOVA with Dunnett’s multiple comparisons test was performed for B-D; *P = <0.05, **P = <0.01

### Slide 3
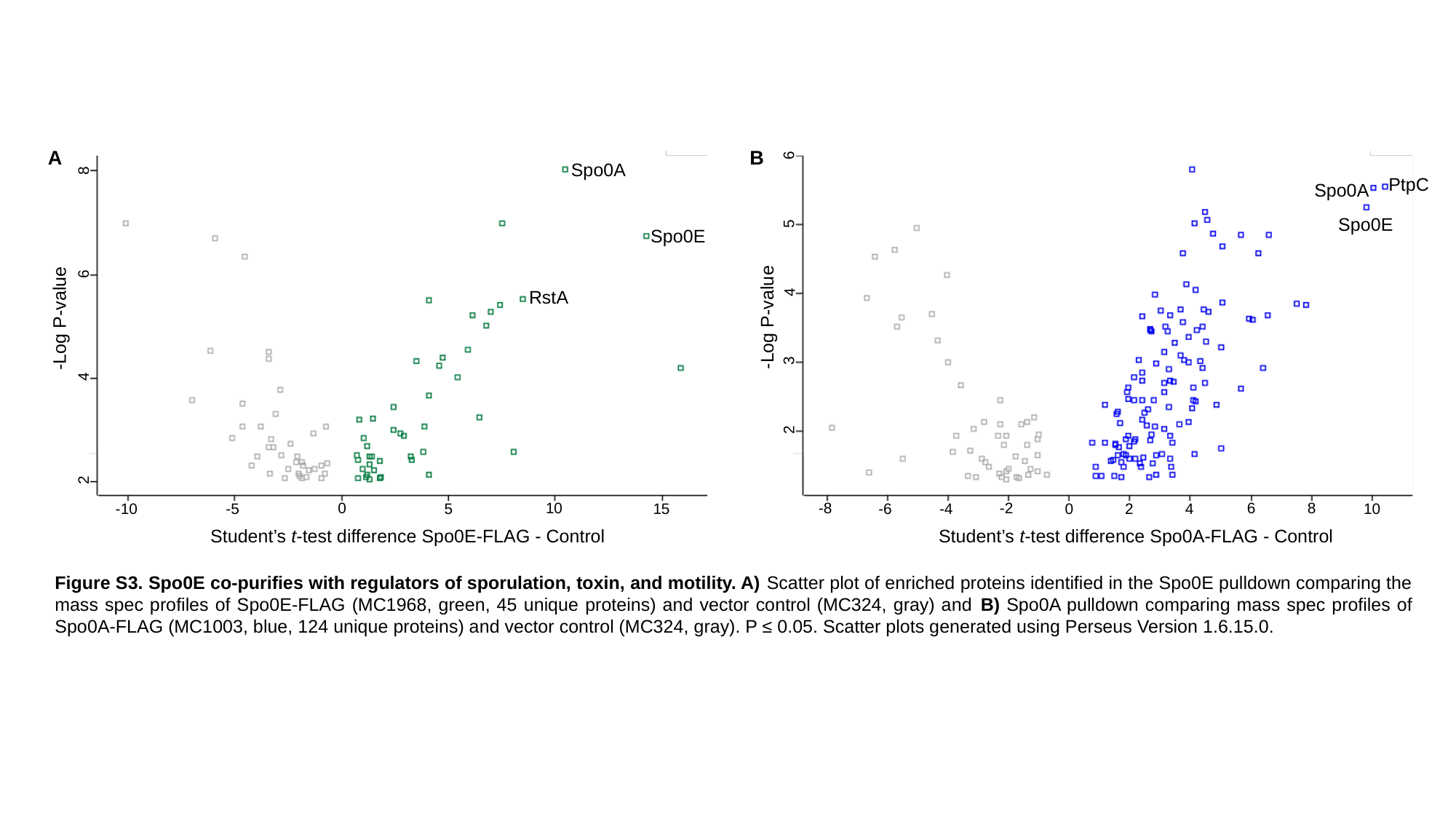

6
A
B
8
Spo0A
PtpC
Spo0A
5
Spo0E
Spo0E
6
4
RstA
-Log P-value
-Log P-value
3
4
2
2
6
8
-2
-8
0
10
-4
4
-6
5
-10
-5
2
10
0
15
Student’s t-test difference Spo0A-FLAG - Control
Student’s t-test difference Spo0E-FLAG - Control
Figure S3. Spo0E co-purifies with regulators of sporulation, toxin, and motility. A) Scatter plot of enriched proteins identified in the Spo0E pulldown comparing the mass spec profiles of Spo0E-FLAG (MC1968, green, 45 unique proteins) and vector control (MC324, gray) and B) Spo0A pulldown comparing mass spec profiles of Spo0A-FLAG (MC1003, blue, 124 unique proteins) and vector control (MC324, gray). P ≤ 0.05. Scatter plots generated using Perseus Version 1.6.15.0.

### Slide 4
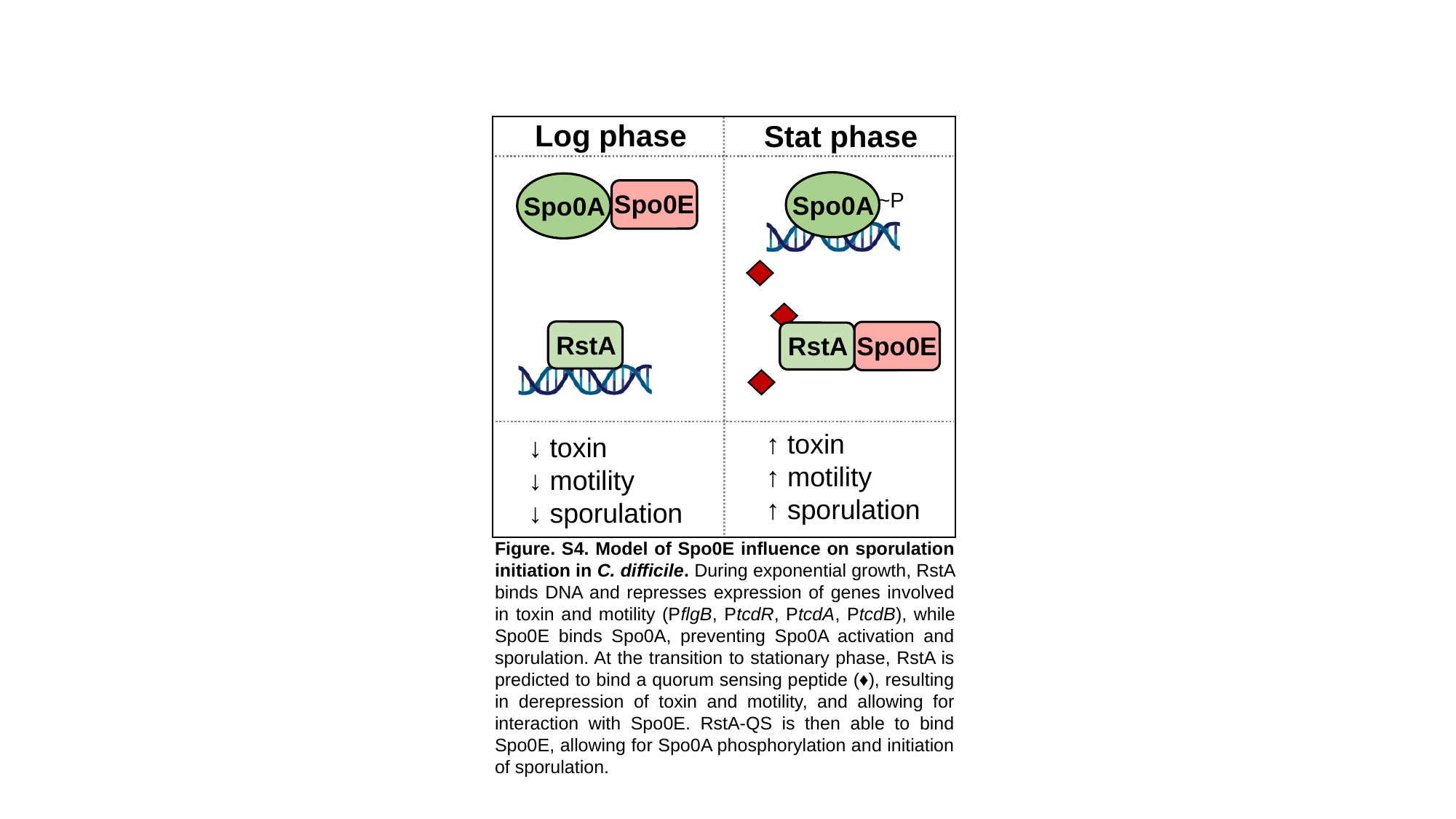

Log phase
Stat phase
Spo0A
~P
Spo0A
Spo0E
RstA
Spo0E
RstA
↑ toxin
↑ motility
↑ sporulation
↓ toxin
↓ motility
↓ sporulation
Figure. S4. Model of Spo0E influence on sporulation initiation in C. difficile. During exponential growth, RstA binds DNA and represses expression of genes involved in toxin and motility (PflgB, PtcdR, PtcdA, PtcdB), while Spo0E binds Spo0A, preventing Spo0A activation and sporulation. At the transition to stationary phase, RstA is predicted to bind a quorum sensing peptide (♦), resulting in derepression of toxin and motility, and allowing for interaction with Spo0E. RstA-QS is then able to bind Spo0E, allowing for Spo0A phosphorylation and initiation of sporulation.
